# Integration of RT-LAMP and Microfluidic Technology for Detection of SARS-CoV-2 in Wastewater as an Advanced Point-of-care Platform

**DOI:** 10.1101/2021.08.18.456880

**Authors:** Ahmed Donia, Muhammad Furqan Shahid, Sammer-ul Hassan, Ramla Shahid, Aftab Ahmad, Aneela Javed, Muhammad Nawaz, Tahir Yaqub, Habib Bokhari

**Author notes:** Correspondence to Habib Bokhari.

## Abstract

Development of lab-on-a-chip (LOC) system based on integration of reverse transcription loop-mediated isothermal amplification (RT-LAMP) and microfluidic technology is expected to speed up SARS-CoV-2 diagnostics allowing early intervention. In the current work, reverse transcriptase quantitative polymerase chain reaction (RT-qPCR) and RT-LAMP assays were performed on extracted RNA of 7 wastewater samples from COVID-19 hotspots. RT□LAMP assay was also performed on wastewater samples without RNA extraction. Current detection of SARS-CoV-2 is mainly by RT-qPCR of ORF (ORF1ab) and N genes so we targeted both to find the best surrogate marker for SARS-CoV-2 detection. We also performed RT-LAMP with/without RNA extraction inside microfluidic device to target both genes. Positivity rates of RT-qPCR and RT-LAMP performed on extracted RNA were 100.0% (7/7) and 85.7% (6/7), respectively. RT-qPCR results revealed that all 7 wastewater samples were positive for N gene (Ct range 37-39), and negative for ORF1ab, suggesting that N gene could be used as a surrogate marker for detection of SARS-CoV-2. RT-LAMP of N and ORF (ORF1a) genes performed on wastewater samples without RNA extraction indicated that all 7 samples remains pink (negative). The color remains pink in all microchannels except microchannels which subjected to RT-LAMP for targeting N region after RNA extraction (yellow color) in 6 out of 7 samples. This study shows that SARS-CoV-2 was successfully detected from wastewater samples using RT-LAMP in microfluidic chips.

## Introduction

Severe acute respiratory syndrome coronavirus 2 (SARS-CoV-2) is a highly transmissible and pathogenic coronavirus and it is the causative agent of the coronavirus disease 2019 (COVID-19) pandemic [1]. Although there are massive coronavirus vaccination campaigns all over the world, strong public health surveillance and rapid diagnostic testing is considered as the best way to control COVID-19 [2-4]. The gold standard to diagnose COVID-19 is reverse transcriptase quantitative polymerase chain reaction (RT-qPCR) [5]. Droplet digital RT-PCR (RT-ddPCR) offers an attractive platform for quantification of SARS-CoV-2 RNA [6]. Factors such as high sensitivity and specificity, requirement of highly trained personnel, and the need of special facilities and high-cost instrumentation limit its application especially in developing countries [7].

Reverse transcription loop-mediated isothermal amplification (RT-LAMP) is an isothermal nucleic acid amplification technique that is being widely used as point-of-care detection of SARS-CoV-2 in clinical samples [8]. RT-LAMP possesses some fundamental advantages such as sensitivity, speed, exclusion of a thermal cycler, and robustness to sample inhibitor making it a promising alternative to RT-qPCR [9]. LAMP takes less than one hour for amplifying the genetic material of the pathogen, and requires a set of four to six primers, ensuring high specificity [10]. Amplification product of LAMP can be confirmed using different procedures such as changes in fluorescence using intercalating dyes, DNA probes with gold nanoparticles [11, 12], changes in turbidity caused by magnesium pyrophosphate precipitate [13], pH indicators, or gel electrophoresis followed by UV detection [14]. The most frequently used method is based on color change of colorimetric master mix containing a visible pH indicator for rapid and easy detection [4, 14, 15].

Although inhalation of aerosol/droplet and person-to-person contact are the major transmission routes of SARS-CoV-2, current evidence points out that the viral RNA is detected in wastewater, urging the need to better understand wastewater as potential source of epidemiological data and human health risks, which can be applied as an early warning system [16-19]. SARS-CoV-2 may cause asymptomatic or pauci-symptomatic infections [20-22], which could add more limitations to determine the actual degree of SARS-CoV-2 circulation in a community. In the meantime, wastewater surveillance can give less biased method of estimating the spread of infection in different places, especially in developing countries, where resources for clinical diagnosis are sparse and limited [23]. Currently, detection of SARS-CoV-2 in wastewater primarily relies on RT-qPCR [24-26], which is laborious, costly, time-consuming, and requires extensive personnel skills [4, 15]. Wastewater-based epidemiology is an alternative method to predict virus spread and it considered as an early warning pandemic through detecting pathogens in wastewater [16, 27]. SARS-CoV-2 biomarkers can be detected in the wastewater and/or sewer system, because the SARS-CoV-2 can be isolated from the infected patients’ urine and feces [28]. Therefore, wastewater analysis in communities is a potential method to track infected people, and to monitor the epidemiology of the communities [29].

The field of microfluidics provides an alternative to the time-consuming bench assays [30]. Micro-electromechanical systems and microelectronics technologies have an important role in the emergence of microfluidic devices, which are able to manipulate minute amounts of fluids and extract information from it, offering the potential to quickly acquire information from the small sample volumes [31].

Traditional RT-qPCR machines are usually bulky and relatively expensive. Besides, the RT-qPCR process is time-consuming [32]. On the other hand, the portable, cheap, and disposable feature of these microfluidic devices makes them suitable for wide range of applications [33]. Another important advantage of microfluidic devices over RT-qPCR is the possibility to integrate RNA extraction inside these devices, which also allows for quick and precise detection of viral RNA [34]. Microfluidics has increasingly been used for point of care testing or bedside. There are many available microfluidic devices for early diagnosis of diseases or other health-related conditions such as pneumonia, glucose level, and pregnancy test by the detection of target elements [30, 35]. In recent years, viruses could be also detected using microfluidic devices [36, 37]. Microfluidic devices promise cheaper, faster, sensitive and easy□to□use methods, so they have a high potential to be an alternative way for the viral RNA detection [38]. Microfluidic devices have previously been applied for detection of RNA viruses such as HIV [39], Hepatitis A virus [40], H1N1 [41], Zika [42], and norovirus [40], with acceptable results.

In the present study, we aim to evaluate the efficacy of RT-LAMP to detect SARS-CoV-2 in wastewater as a quick method to provide early warning system for COVID-19 transmission in the community. Current detection of SARS-CoV-2 is mainly by RT-qPCR of ORF (ORF1ab) and N genes so we try to find the best surrogate gene marker for SARS-CoV-2 detection. We also aim to assess the application RT-LAMP in microfluidic device as an advanced method to detect SARS-CoV-2 in wastewater.

## Material and methods

### Wastewater sampling

Grab sampling technique was used to collect untreated wastewater samples (sewage samples). During the peak morning flow, 500 ml of wastewater was collected from the midstream into a leak proof, sterile container at a downstream sampling site. Seven wastewater samples were collected in early morning from hot COVID-19 spots in Islamabad, capital of Pakistan on 4 April 2021 and were kept at 4 °C. In the following day (5 April 2021) early morning, samples were transported at 4 °C to the BSL-3 facility at the Institute of Microbiology (IM), University of Veterinary and Animal Sciences (UVAS) Lahore, Pakistan. All experiments in this study were performed in triplicate (three times).

### Sample processing and RNA extraction

RNA of each wastewater sample was extracted in BSL-3 of IM, UVAS Lahore, Pakistan. Before extraction, each sample was vortexed thoroughly and 1 ml of the sample was transferred to microfuge tube. Samples were centrifuged at 5000 rpm for 15 minutes at 4 °C. The supernatant was used for RNA extraction [43]. RNA was extracted using Viral Nucleic Acid Extraction Kit II Geneaid (Geneaid Biotech, Taiwan), according to the manufacturer’s protocol. The RNA was stored at -80°C, and used as a template for both RT-qPCR, and RT□LAMP. We performed RNA extraction step on the seven samples directly without virus concentration step to check if virus concentration step is necessary or can be skipped in RT-LAMP of wastewater samples.

### RT-qPCR analysis

RT-qPCR analysis of the seven wastewater samples was performed by using the commercially available kit (2019-nCoV Nucleic Acid Diagnostic Kit, Sansure Biotech Inc., China). This kit is used for detection of the ORF (ORF1ab) and N genes of SARS-CoV-2. According to Sansure protocol, we selected FAM (ORF-1ab region) and ROX (N gene) channels. Each reaction mixture contained 26μl of 2019-nCoV-PCR Mix, 4μl of 2019-nCoV-PCR Enzyme Mix, and 20 μl RNA extract so the final volume will be 50 μl. Thermal cycling reactions are shown in supplementary Table 1. RT-qPCR analysis was run on CFX96 real-time thermal cycler (Bio-Rad, USA). All RT-qPCR reactions also had positive and negative controls.

For all RT-qPCR, and RT-LAMP assays, we used positive controls and negative controls provided from Sansure kit. According to Sansure kit, the positive control contains *in vitro* transcriptional RNA for ORF1ab, N gene and internal control RNase P gene. The negative control in Sansure kit contains normal saline. Interpretation of RT-qPCR results were shown in supplementary Table 2.

### RT□LAMP assays performed with/without RNA extraction

To detect SARS-CoV-2 RNA with RT-LAMP, we used the WarmStart Colorimetric LAMP 2X Master Mix (DNA and RNA) from New England Biolabs (USA), which contains two enzymes, an engineered reverse transcriptase (RTx) and a warmStart strand-displacing DNA polymerase (Bst 2.0) in a special low-buffer reaction solution containing a visible pH indicator for easy and rapid detection of LAMP (DNA) and RT-LAMP (RNA) reactions. As a way to avoid nonspecific priming reactions, there are oligonucleotide-based aptamers in the reaction mixture to work as reversible temperature-dependent inhibitors, ensuring that the reaction only starts at high temperature (WarmStart) (https://international.neb.com/products/m1800-warmstart-colorimetric-lamp-2x-master-mix-dna-rna). This aim of this system is to provide a fast, clear visual detection of amplification based on the production of protons, leading to pH drop due to extensive DNA polymerase activity in a LAMP reaction, producing a color change from negative pink (alkaline) to positive yellow (acidic).

RT□LAMP assays were performed on the seven wastewater samples according to Zhang *et al*. [44] with two sets of LAMP primers targeting ORF (ORF1a) and N genes [44] (supplementary Table 3). We selected these two sets, because Zhang *et al*. [44] concluded that these two sets were the best performing among five tested sets targeted the ORF (ORF1a) and N genes [44]. The 5′ frameshift nature of polyproteins (ORF1a/ORF1ab) [45] allows us to use primers for ORF1a, ORF1ab genes in RT-LAMP, RT-qPCR respectively. In other way, ORF1ab, the largest gene, contains overlapping open reading frames that encode polyproteins PP1a and PP1ab [46] so ORF1a is part of ORF1ab.

In brief, the assay was performed in a 20 μl reaction mixture containing 2 μL of 10x primer mix of 16 μM (each) of Forward Inner Primer (FIP) and Backward Inner Primer (BIP), 2 μM (each) of F3 and B3 primers, 4 μM (each) of Forward Loop (LF) and Backward Loop (LB) primers, 10 μL of WarmStart Colorimetric Lamp 2X Master Mix (M1800) (New England Biolabs, USA), 5 μL of DNAse, RNAase free water (Invitrogen, USA), and 3 μl of RNA template. The reaction mixture was set at 65 °C for 30 minutes on a pre-heated dry bath. Yellow color indicates positive reaction, where pink indicates negative one. We also performed RT□LAMP on the 7 samples directly without RNA extraction to check if RNA extraction is necessary or can be omitted in RT-LAMP of wastewater samples.

### RT□LAMP assays in microfluidic device

The microchips were designed using CAD software (SolidWorks, Dassault Systemes) and 8 mm long microchannels were micromachined on a polymethyl methacrylate (PMMA) piece (1.2 mm thickness) with a cross-section of 0.6 × 0.6 mm (width x depth). A sample container was also micromilled using 5 mm thick PMMA sheets to load 10 μL sample into each well. Before use, microcapillaries were coated with polyvinyl alcohol (PVA) to convert their surfaces from hydrophobic to hydrophilic, allowing liquid to rise in the microchannels [47, 48] with modifications. In brief, 2% PVA (molecular weight 146000–186000) was prepared in deionized water (DI) water and used to fill the microchannels at room temperature. After 30 minutes, excess PVA was removed and the microchannels were dried by injection of compressed air, and placed in an oven for 15 min. Then, the microchip was simply dipped into the RT-LAMP reaction solution loaded via capillary action without need of pump. The microchips were set at 65 °C for 30 minutes on a pre-heated dry bath. We performed RT-LAMP inside microfluidic device to target ORF (ORF1a), and N genes of the first sample. We used both RNA extract and direct wastewater sample (without RNA extraction). RT-LAMP was performed on the microfluidic device using the same conditions as previously described.

### Statistical analyses

Statistical analyses were conducted using graphpad prism. Statistical significance was assessed using the Student T test.

## Results

### Analysis of RT-qPCR and RT-LAMP done on RNA extracts

The results of 7 samples tested by RT-LAMP and RT-qPCR are shown in Table 1. Positivity rates of RT-qPCR and RT-LAMP were 100.0% (7/7) and 85.7% (6/7), respectively. RT-qPCR results revealed that all 7 wastewater samples were positive for N gene (Ct range 37-39), and negative for ORF (ORF1ab). All samples had Ct for internal control (CY5 channel) less than 40. Therefore, according to the guidelines of RT-qPCR kit manufacturer company (Sansure), all 7 samples were positive for SARS-CoV-2.

**Table 1:** Comparison of detection accuracy between RT-LAMP and RT-qPCR.

| Sample ID | RT-qPCR |  | RT-LAMP |  |
| --- | --- | --- | --- | --- |
|  | ORF (ORF1ab) | N | ORF (ORF1a) | N |
| 1 | 41 | 39 | - | + |
| 2 | N/A | 38 | - | + |
| 3 | 42 | 37 | - | - |
| 4 | N/A | 39 | - | + |
| 5 | 41 | 35 | - | + |
| 6 | N/A | 38 | - | + |
| 7 | 44 | 37 | - | + |

+ Positive reaction; - Negative reaction; N/A not detected

Of 7 wastewater samples positive for SARS-CoV-2 by RT-qPCR, 6 were positive by RT-LAMP (sensitivity of 85.7%). As shown in Figure 1, results of RT-LAMP targeted N gene of SARS-CoV-2 revealed that 6 samples (sample id 1, 2, 4, 5, 6, and 7) showed color change to yellow color. As shown in Figure 2, results of RT-LAMP targeted ORF (ORF1a gene) of SARS-CoV-2 revealed that all 7 samples were negative for ORF (ORF1a gene) with pink color.

**Figure 1:**
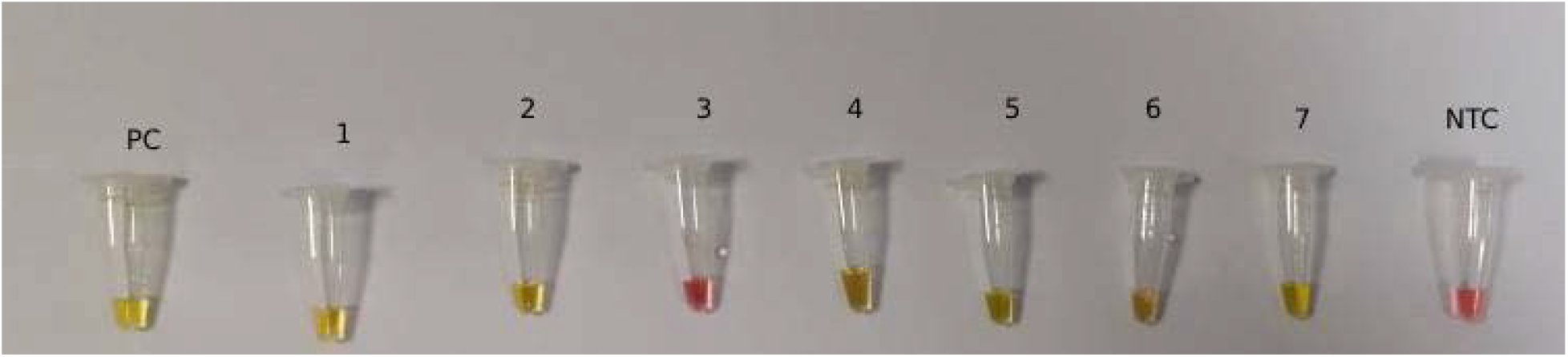
The colorimetric detection of SARS-CoV-2 (N gene) using RT-LAMP of the seven wastewater samples after RNA extraction.

**Figure 2:**
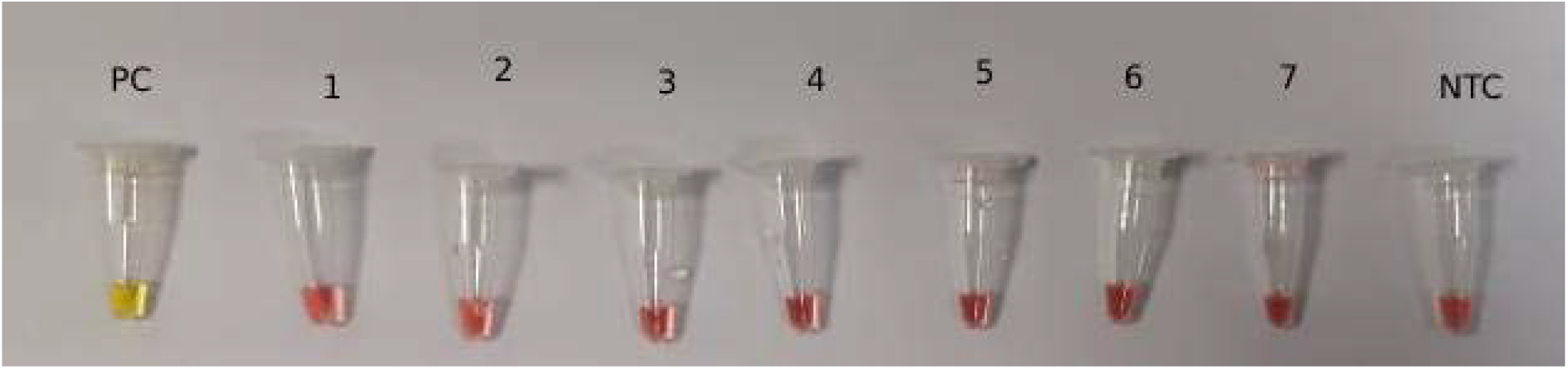
The colorimetric detection of SARS-CoV-2 (ORF1a gene) using RT-LAMP of the seven wastewater samples after RNA extraction.

### RT-LAMP without RNA extraction

We performed RT-LAMP for amplification of N and ORF (ORF1a) genes directly on wastewater samples without RNA extraction, and found that all seven samples remains pink indicating negative results for both N and ORF (ORF1a) genes (Figure 3).

**Figure 3:**
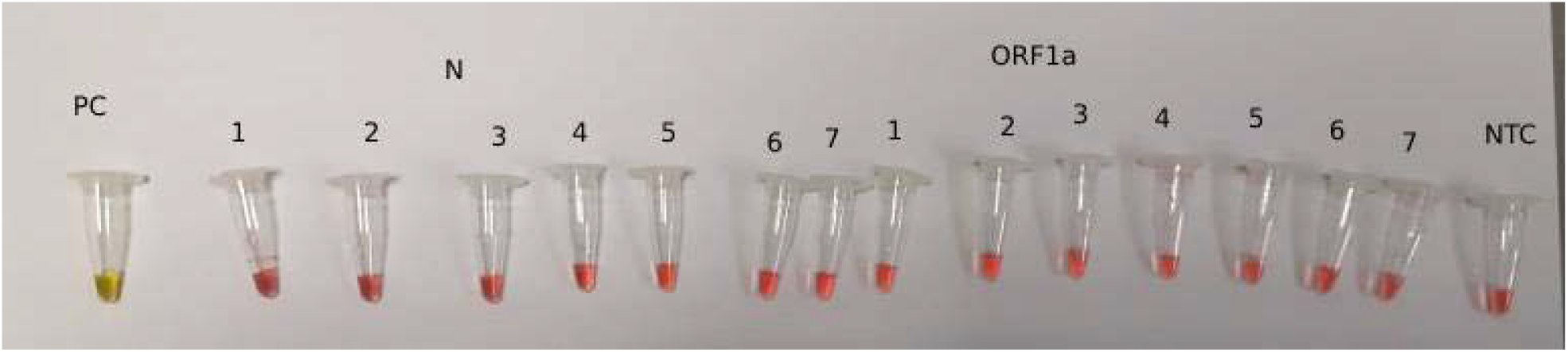
The colorimetric detection of SARS-CoV-2 (N and ORF1a genes) using RT-LAMP of the seven wastewater samples directly without RNA extraction (All seven samples remains pink indicating negative results for both N and ORF1a genes).

### RT□LAMP assays in microfluidic device

Figure 4 shows microfluidic device and wells. Successful loading of RT-LAMP mixtures into microfluidic device is shown in Figure 5. As shown in Figure 6, the color remains pink in all microchannels except the one which subjected to RT-LAMP for targeting N region after RNA extraction (yellow color) in 6 out of 7 samples.

**Figure 4:**
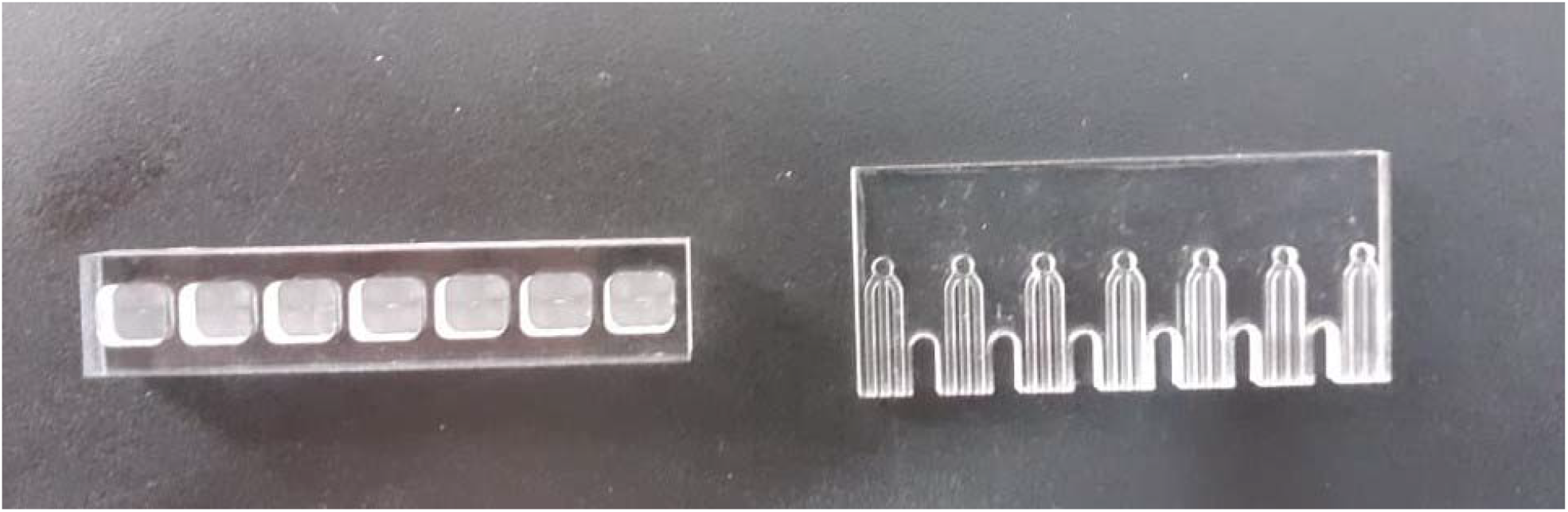
Microfluidic device and wells.

**Figure 5:**
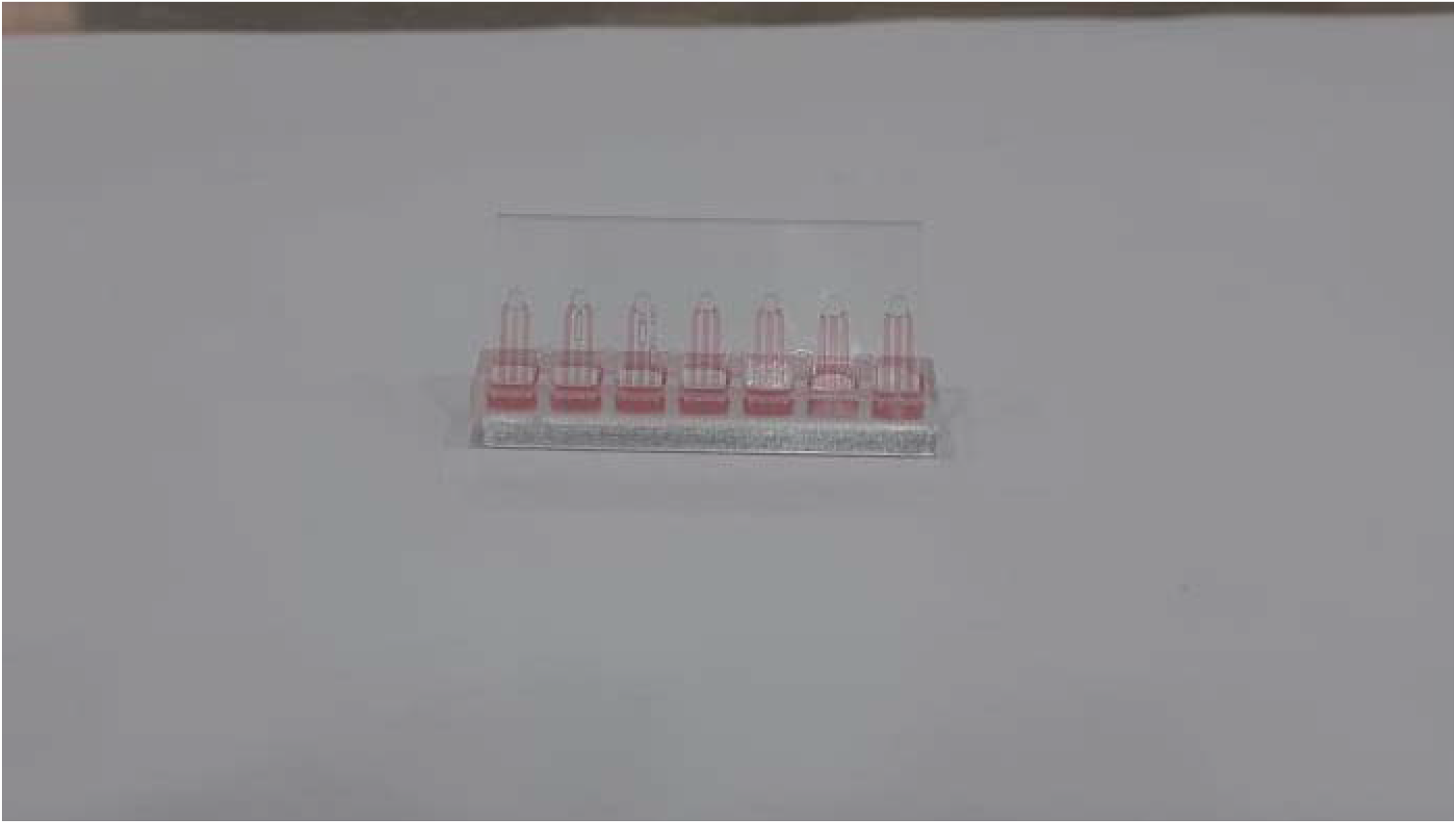
Successful loading of RT-LAMP mixtures into microfluidic device

**Figure 6:**
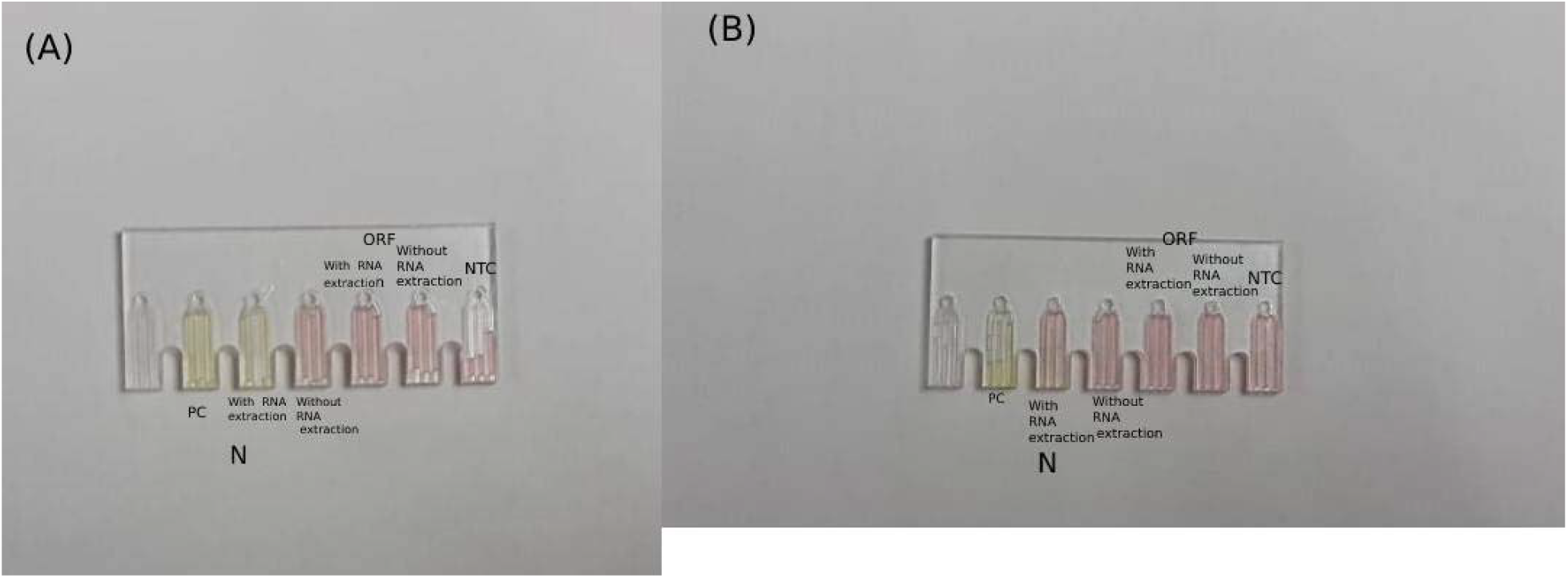
(A) Samples 1, 2, 4, 5, 6, and 7 showed color change to yellow in microchannels of N gene after RNA extraction. (B) Sample 3 did not show color change (remains red even in microchannels of N gene after RNA extraction).

### Statistical analysis

With p value of 0.0001, there is a highly significant difference of SARS-CoV-2 detection with and without RNA extraction (Figure 7). There is no significant difference (P value = 0.6) between RT-qPCR and RT-LAMP assays used for detection of SARS-CoV-2 (Figure 8). There is a highly significant difference of detection of ORF and N genes of SARS-CoV-2 (P value =0.0001) (Figure 9).

**Figure 7:**
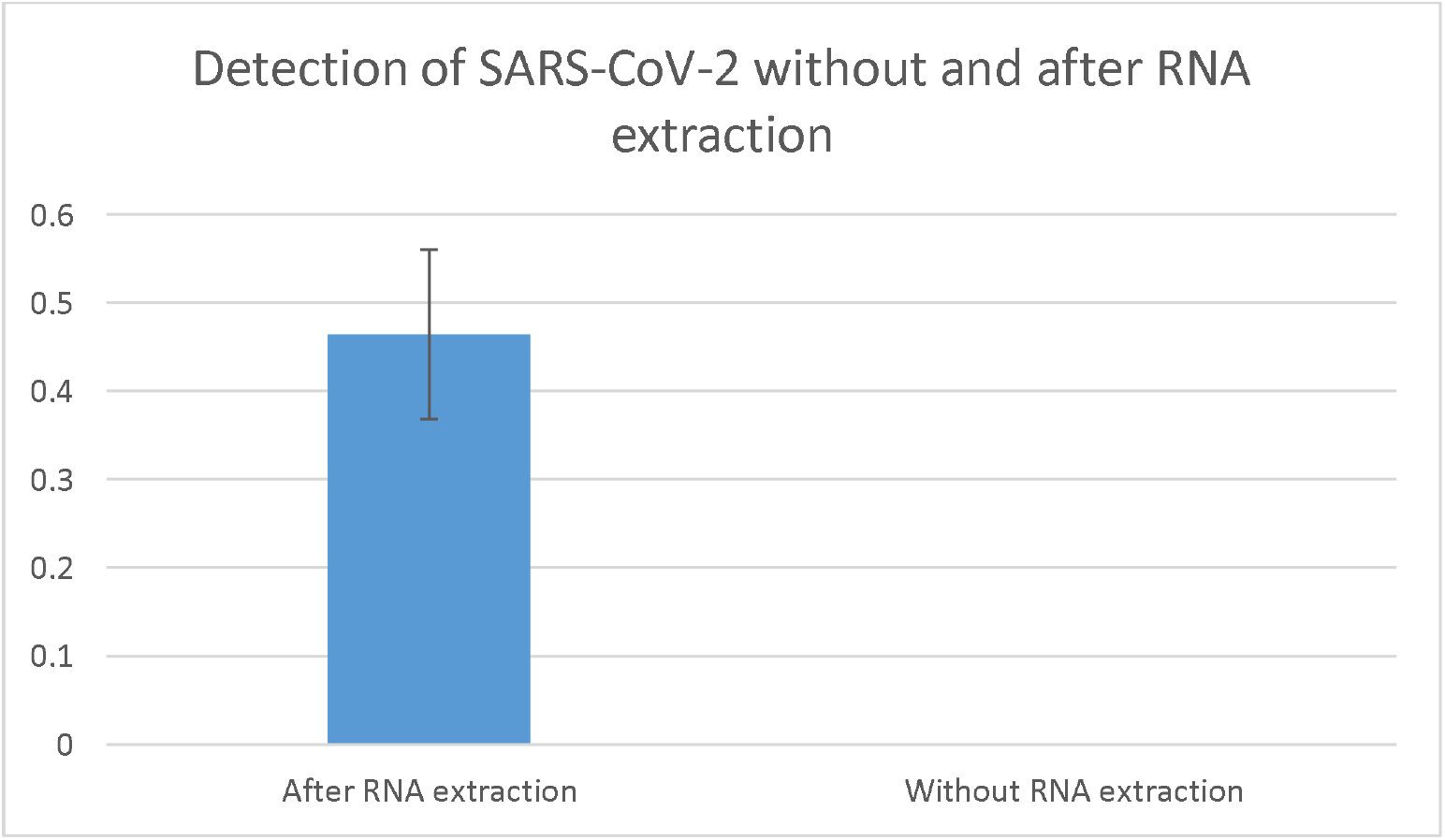
Detection of SARS-CoV-2 without and after RNA extraction (P value =0.0001 highly significant).

**Figure 8:**
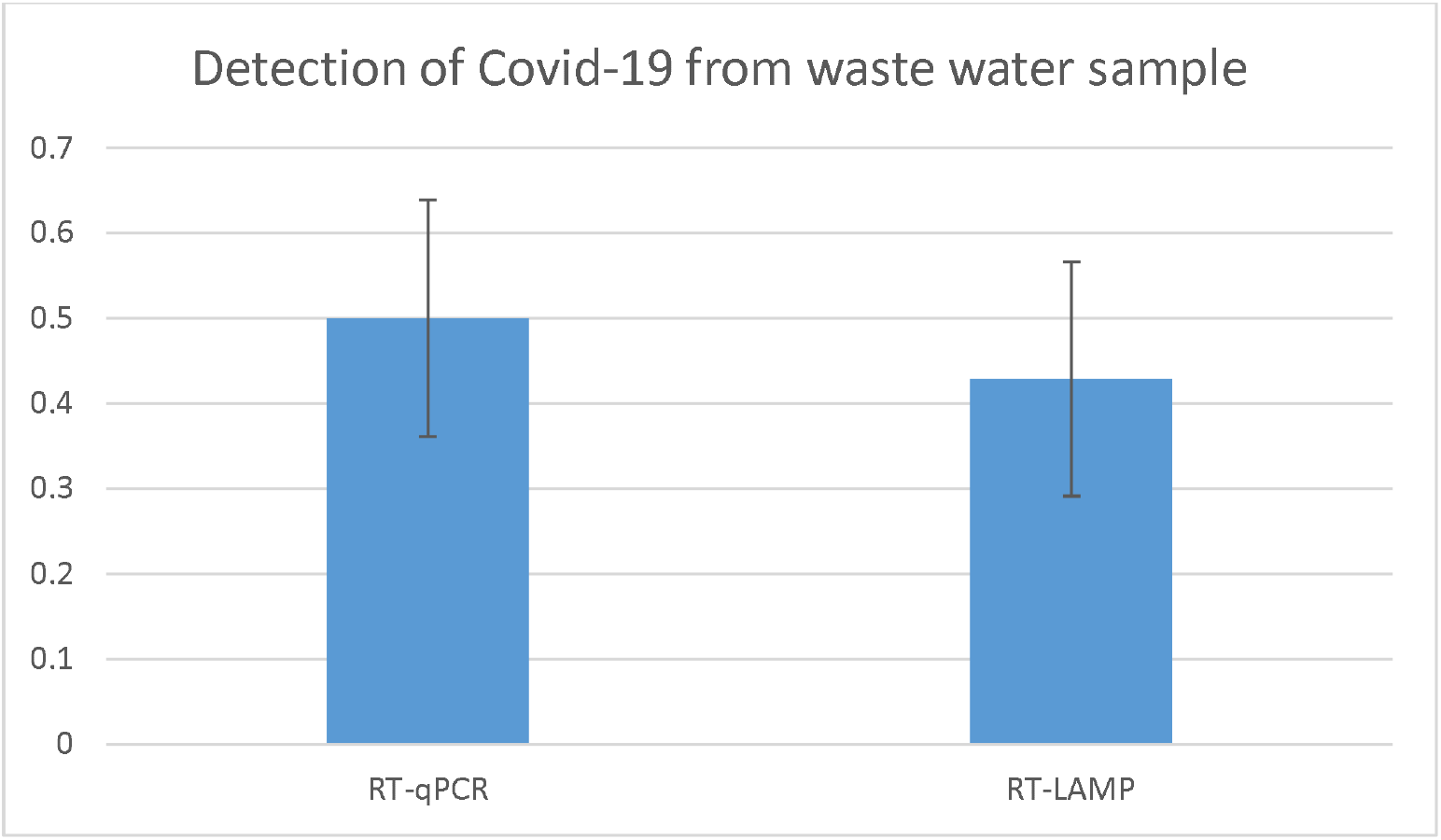
Detection of SARS-CoV-2 using RT-qPCR and RT-LAMP (P value = 0.6 non-significant).

**Figure 9:**
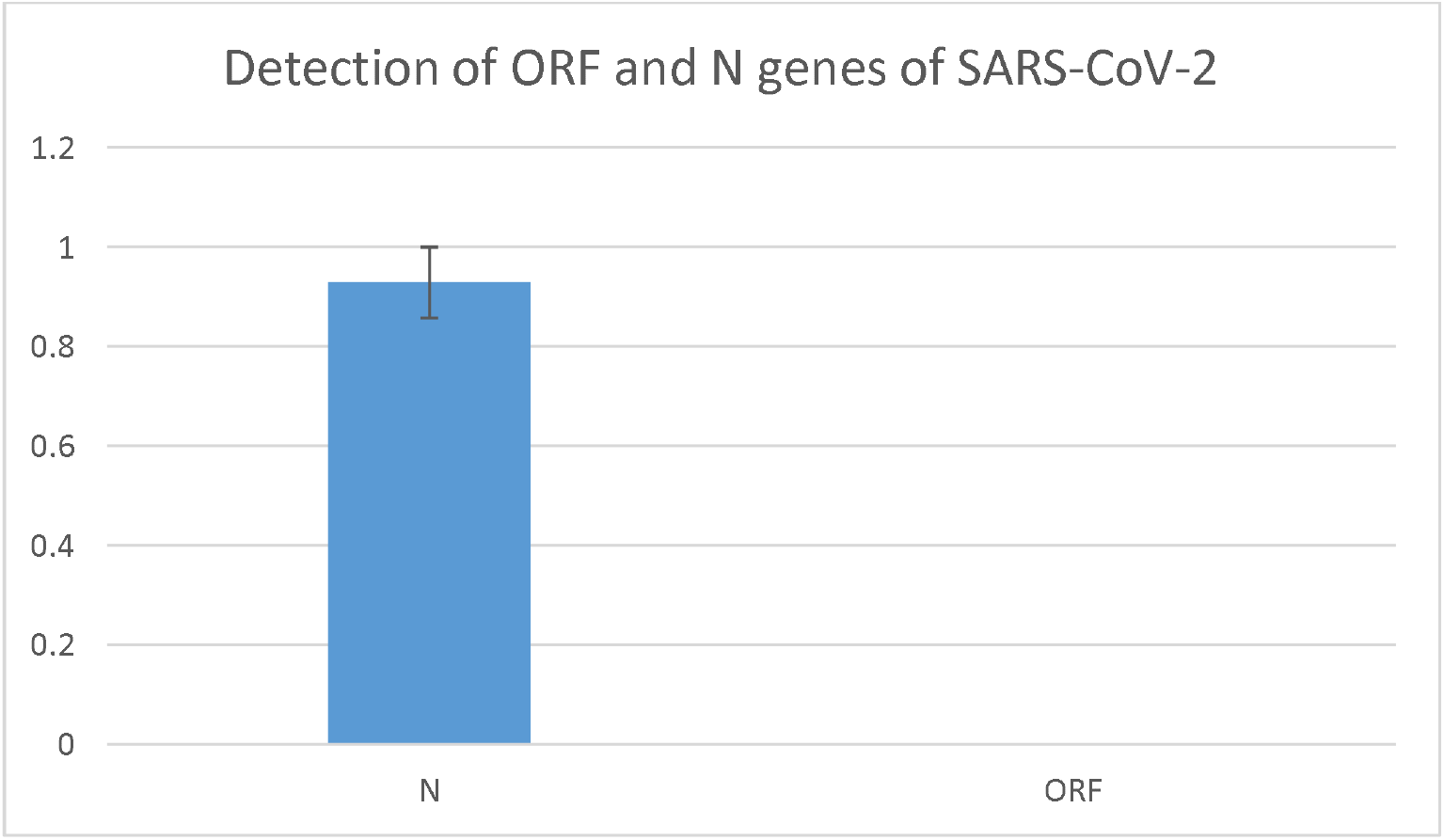
Detection of ORF and N genes of SARS-CoV-2 (P value =0.0001 highly significant)

## Discussion

RT-LAMP recently emerged as an alternative point-of-care test for detection of SARS-CoV-2, including clinical sample testing, with reaction time varying between 20 and 40 minutes [44, 49, 50]. RT-LAMP has some fundamental advantages such as constant temperature amplification, elimination of a thermal cycler, quick test result, and wide diagnostic capacity, while keeping similar specificity and sensitivity, thus making it more appropriate than the RT-qPCR for monitoring a pandemic such as COVID-19 [51]. Most of published papers are about using of RT-LAMP for detection of SARS-CoV-2 in patient or clinical samples [14, 50, 52-56], with few of them have focused on detection of SARS-CoV-2 using RT-LAMP in wastewater or sewage samples [57, 58]. Bivins and colleagues used tampons as passive swabs for sample collection and RT-LAMP to detect SARS-CoV-2 RNA in wastewater, and they found that the workflow were available within three hours of sample collection [59]. SARS-CoV-2 RdRp-based LAMP assay detected the virus RNA in 26/28 (93%) of RT-PCR positive COVID-19 clinical samples with 100% specificity (n = 7) within 20 min [57]. We found that results of RT-LAMP for these wastewater samples were largely consistent with those of RT-qPCR, with 6 out of 7 (85.7%) samples were positive for both RT-LAMP and RT-qPCR. Only one sample (sample id 3) tested positive by RT-qPCR was negative by our RT-LAMP.

The Nucleocapsid protein can regulate the replication, transcription and packaging, and it is important for viral viability. There is a growing interest in studying the N protein for vaccine development because of its highly immunogenic and its highly conserved amino acid sequence [60]. Currently, the detection of SARS-CoV-2 RNA is mainly performed by RT-qPCR detection of two target genes, including ORF1ab and N [61]. We found that the Ct values of N gene for all of our 7 wastewater samples were lower than the Ct values of ORF1ab gene, suggesting that N gene is the most important gene when monitoring SARS-CoV-2. Our results were similar to what previously reported [62], where the highest proportion of positive results among COVID-19 was the N gene, followed by both ORF1ab and N. They found that the main positive fragment is the N gene, and the proportion of those positive for single ORF1ab was very low [62]. Therefore, we recommended monitoring of the N gene as surrogate marker for detection in wastewater samples, reducing the time and cost of nucleic acid detection. Loying and colleagues [63] studied the dynamics of ORF1ab and N genes among hospitalized COVID-19 positive patients, and they found that the persistent of positivity of N gene is significantly for more duration than ORF1ab, indicating that N gene requires longer duration of days to become negative than ORF1ab. This also underscores our proposition that N gene should be considered as a surrogate marker for detection in clinical and wastewater samples.

One of the most important advantage of RT-LAMP is its ability to detect SARS-CoV-2 directly from clinical samples without the need of RNA extraction [52, 64]. Wei and colleagues [64] developed and tested a highly sensitive and robust method based on RT-LAMP that uses readily available reagents and a simple heat block using contrived spike-in and actual clinical samples. They directly tested clinical nasopharyngeal swab samples in viral transport media without previous time-consuming and laborious RNA extraction with results in just 30 min. Mautner and colleagues [52] developed RT-LAMP assay to directly detect SARS-CoV-2 from pharyngeal swab samples without previous RNA extraction. They found that this method is 10 times cheaper and 12 times faster than RT-qPCR, depending on the assay used. Previous study performed RT-LAMP on wastewater samples after RNA extraction and virus concentration [57]. According to CDC guidelines for wastewater surveillance testing methods (https://www.cdc.gov/healthywater/surveillance/wastewater-surveillance/testing-methods.html), small volumes of wastewater (e.g., 1 ml) may be tested without additional concentration processes if levels of SARS-CoV-2 RNA are sufficiently high in wastewater. Therefore, we targeted hotspots in Islamabad with recorded high COVID-19 cases, and took 7 wastewater samples to see the possibility to detect SARS-CoV-2 in wastewater without RNA extraction and virus concentration using direct RT-LAMP alone. Although RNA extraction may be omitted in RT-LAMP performed on clinical samples [52, 64], we found that RNA extraction is necessary in RT-LAMP performed on wastewater samples. However, Ongerth and Danielson detected SARS-CoV-2 in raw sewage samples with no preliminary sample processing for virus concentration and RNA extraction [65].

Microfluidic techniques are emerging as disposable and cost-efficient tools for rapid diagnosis of viral infection [37]. Since microfluidic devices are sensitive, cheaper, faster, and easy□to□use methods, they have a high potential to be an alternative way for the viral RNA detection [38]. Safavieh *et al*. developed RT□LAMP cellulose□based paper microchips and amplified the target RNA using the RT□LAMP technique and detected the HIV□1 virus via the electrical sensing of LAMP amplicons [39]. Fraisse *et al*. designed RT□PCR integrated microfluidic device to detect Hepatitis A and noroviruses in the gut [40]. Song *et al*. developed RT□LAMP integrated microfluidic for detection of Zika virus [42]. Recently, Kim *et al*. designed RT□PCR integrated microfluidic device for detecting of H1N1 influenza in saliva [41].

We used RT-LAMP in microfluidic chip for detection of SARS-CoV-2 in wastewater. After coating of microfluidic chip with PVA, RT-LAMP mixture can be successfully uploaded into microchannels. Then, we observed color change in microfluidic chip after placing it in pre-heated dry bath at 65 °C for 30 minutes. Although we found that detection of SARS-CoV-2 in wastewater using RT-LAMP in microfluidic chip requires RNA extraction, we propose that our workflow (without RNA extraction) could work with clinical samples since there are many reports about successful detection of SARS-CoV-2 in clinical samples using RT-LAMP without RNA extraction [52, 64, 66, 67]. If RNA extraction could be achieved in this microfluidic chip, this could greatly improve the results of this chip. Mauk and colleagues developed simple plastic microfluidic chip for nucleic acid-based testing of blood, other clinical specimens, water, food, and environmental samples [34]. They combines isolation of nucleic acid by solid-phase extraction; isothermal enzymatic amplification such as LAMP, nucleic acid sequence based amplification, and recombinase polymerase amplification; and real-time optical detection of DNA or RNA analytes. Recent study combined two-stage isothermal amplification assay into the integrated microfluidic platform to detect SARS-CoV-2 and human enteric pathogens in wastewater within one hour [68]. Although we could not detect clear color change in sample not subjected to RNA extraction, we propose that performing RNA extraction inside microchannels could greatly improve results.

Rodriguez-Mateos and colleagues integrated on-chip platform coupling RNA extraction based on immiscible filtration assisted by surface tension, with RNA amplification and detection via colorimetric RT-LAMP, using ORF1a and N genes of SARS-CoV-2 [69]. They performed on-chip processes for the SARS-CoV-2 spiked water samples, but not on wastewater samples. In our study, we successfully integrated microfluidic technology and RT-LAMP on wastewater samples from COVID-19 hotspots, which could give better idea about spread of the SARS-CoV-2 in the community. Our work is expected to pave the road for designing readymade microfluidic chip with specific chamber for RNA extraction then another amplification chamber for RT-LAMP. This amplification product could be also labeled with a fluorophore reporter that could be excited with a LED light source and monitored in situ in real time with a photodiode or a CCD detector (such as available in a smartphone).

## Conclusion

RT-LAMP has been emerging as a great alternative to the RT-qPCR because RT-LAMP is a specific, sensitive, fast, cheap, and easy□to□use method. We successfully detected SARS-CoV-2 through color change (yellow color) in our positive wastewater samples having Ct >30. We also found that the Ct values of N gene for all of our wastewater samples were lower than the Ct values of ORF1ab gene, suggesting that N gene could be used as a surrogate marker for monitoring and surveillance of environmental circulating SARS-CoV-2. We successfully detected SARS-CoV-2 from wastewater samples using RT-LAMP in microfluidic chips. This will provide an opportunity for developing more robust and economical approach for using this microchip device along with RT-LAMP as an advanced method for detection of SARS-CoV-2 in different samples.

## Conflict of interest

The authors report no conflicts of interest in this work.

## Availability of data and material (data transparency)

Data available within the article or its supplementary materials

## Code availability (software application or custom code)

Not applicable

## Acknowledgments

This paper is part of Ph.D. research work of the first author (Ahmed Donia) who is extremely thankful to Professor Habib Bokhari for supervision of this work. He would like to warmly thank all staff at The Biosafety Level-3 Laboratory for Emerging Pathogens, Institute of Microbiology, University of Veterinary and Animal Sciences (UVAS), Lahore, Pakistan for extreme support to carry out this work. He would like also to warmly thank The World Academy of Science (TWAS) and Microbiology and Public Health Laboratory, COMSATS University Islamabad (CUI) Pakistan, and Kohsar University Murree.

## Author contributions

AD and HB conceived and designed the study. AD, MFS, RS coordinated, carried out the experiments, and analyzed the data. AD drafted the original manuscript. AD, MFS, RS, AA, AJ, MN, TY, and HB did necessary editing of the manuscript. All authors read and approved the manuscript.

## Funding

This research received no external funding

## Notes

### Competing Interest Statement

The authors have declared no competing interest.

