## Supplemental Tables for "Integration of RT-LAMP and Microfluidic Technology for Detection of SARS-CoV-2 in Wastewater as an Advanced Point-of-care Platform"

**Supplementary Table 1: Thermal cycling reactions of RT-qPCR of SARS-CoV-2 according to Sansure protocol.**

|  | Steps | Temperature. | Time. | Cycle No. |
| --- | --- | --- | --- | --- |
| 1 | Reverse transcription | 50°C | 30 min. | 1 |
| 2 | cDNA predenaturation | 95°C | 1 min. | 1 |
| 3 | Denaturation | 95°C | 15 sec. | 45 |
|  | Annealing, extension and fluorescence collection | 60°C | 30 sec. |  |
| 4 | Device cooling | 25°C | 10 sec. | 1 |

**Supplementary Table 2: Explanation of detection result according to Sansure protocol.**

| Conclusion | Amplification results |
| --- | --- |
| SARS-CoV-2 Positive | There is typical S-shape amplification curve detected at FAM and/or ROX channel, and the amplification curve which is detected at CY5 (internal control) channel, Ct≤40. |
| SARS-CoV-2 Negative | There is no typical S-shape amplification curve (No Ct) or Ct＞40 detected at FAM and ROX channel, and the amplification curve which is detected at CY5 channel (internal control), Ct ≤ 40. |

**Supplementary Table 3: Sequences of amplicons and LAMP primers**

| **LAMP primer** | **Sequence** |
| --- | --- |
| **ORF1a** |  |
| ORF1a - F3 | CTGCACCTCATGGTCATGTT |
| ORF1a -B3 | AGCTCGTCGCCTAAGTCAA |
| ORF1a -FIP | GAGGGACAAGGACACCAAGTGTATGGTTGAGCTGGTAGCAGA |
| ORF1a -BIP | CCAGTGGCTTACCGCAAGGTTTTAGATCGGCGCCGTAAC |
| ORF1a -LF | CCGTACTGAATGCCTTCGAGT |
| ORF1a -LB | TTCGTAAGAACGGTAATAAAGGAGC |
| **N** |  |
| N-F3 | TGGCTACTACCGAAGAGCT |
| N-B3 | TGCAGCATTGTTAGCAGGAT |
| N-FIP | TCTGGCCCAGTTCCTAGGTAGTCCAGACGAATTCGTGGTGG |
| N-BIP | AGACGGCATCATATGGGTTGCACGGGTGCCAATGTGATCT |
| N-LF | GGACTGAGATCTTTCATTTTACCGT |
| N-LB | ACTGAGGGAGCCTTGAATACA |
